# Genomic Foundation Models Reveal Chromatin-Domain-Scale Transposable Element Impacts on Rice Genome Architecture

**DOI:** 10.64898/2026.05.11.724192

**Authors:** Jingchao Fan, Hui Zhao, Qianjie Lv, Xiaoli Wang, Rui Man, Nengfu Xie, Zhonghe Zhao

## Abstract

Alignment-based detection of transposable element (TE) insertion polymorphisms suffers from reference bias and multi-mapping errors in repetitive genomic regions, creating a fundamental validation bottleneck for population-scale structural variant catalogs. Here, we demonstrate that the OneGenome-Rice (OGR) genomic foundation model (GFM)—a 1.25 billion parameter Mixtral architecture trained on 422 rice genomes without TE annotations—provides an entirely orthogonal, alignment-free approach that resolves TE-mediated structural divergence at chromatin-domain resolution. At the CTB4a cold-tolerance locus on chromosome 4, OGR embeddings revealed that the aus subpopulation (NONA_BOKRA) carries 2.2-fold higher structural divergence from indica than japonica, consistent with its 728 subpopulation-exclusive cold-protective TE insertions. Sliding-window analysis across 4.4 megabases identified a 25.6-fold divergence enhancement at TE clusters relative to the conserved CTB4a gene body. Critically, the minimal effective resolution was established at approximately 20 kilobases—corresponding to the median size of topologically associating domains (TADs) in the rice genome—while individual TE sites at 500 base pairs were undetectable (P = 0.94). Non-neural baselines confirmed the signal derives from learned representations of genomic context rather than simple nucleotide statistics. These findings establish GFMs as orthogonal validation tools for population-scale TE genotyping and provide computational evidence that TE functional effects are organized at the chromatin-domain level, with direct implications for prioritizing functional TE variants in crop breeding.

## Introduction

Transposable elements (TEs) are the single largest component of plant genomes, constituting 45–55% of the rice (Oryza sativa L.) genome and over 85% of the maize genome (International Rice Genome Sequencing Project, 2005; Schnable et al., 2009; Song et al., 2021). Far from being inert ‘junk DNA,’ TEs are increasingly recognized as dynamic contributors to genome evolution, gene regulation, and environmental adaptation (Lisch, 2013; Bennetzen and Wang, 2014). The mobilization of TEs—through retrotransposition and DNA transposition—generates insertion polymorphisms that can alter gene expression, create novel regulatory elements, and reshape local chromatin architecture (Feschotte, 2008; Chuong et al., 2017). In crops, TE insertions have been causally linked to agronomic traits including grape berry color (Kobayashi et al., 2004), tomato fruit shape (Xiao et al., 2008), and rice stress tolerance (Qian et al., 2025).

The discovery that plant genomes are predominantly composed of TEs was enabled by reference genome assemblies. However, these same reference genomes create a paradox for TE research: the repetitive nature of TE sequences makes them the most difficult genomic features to accurately assemble, genotype, and validate (Goerner-Potvin and Bourque, 2018). All existing TE insertion polymorphism (TIP) detection methods— whether based on split-read alignment, discordant paired-end mapping, or assembly-based approaches—share a fundamental dependency on read alignment to a linear reference genome (Ewing, 2015; Carpentier et al., 2019). This creates two well-documented vulnerabilities: reference bias, where structurally divergent reads fail to align causing false-negative TE calls, and multi-mapping, where conserved TE sequences align to paralogous copies causing ambiguous assignments (Li and Durbin, 2009; Treangen and Salzberg, 2012). In practice, these biases are severe: in pericentromeric regions where TE density exceeds 80%, false-negative rates for alignment-based TE detection can exceed 50% (Zhou et al., 2020; Song et al., 2021), and multi-mapping rates for LTR retrotransposon families like Gypsy and Copia routinely exceed 30% even with paired-end strategies (Carpentier et al., 2019). These challenges are particularly acute in the very genomic territories where TE functional impact may be greatest—centromeric and peri-centromeric regions harbor the highest TE densities but the lowest alignment confidence.

Traditional cross-validation of TE genotyping requires PCR amplification or long-read sequencing (Sudmant et al., 2015), both expensive methods that cannot be applied retroactively to existing population-scale datasets. An orthogonal computational method—one that validates TE calls without sharing the assumptions of alignment-based detection—would substantially enhance confidence in TE polymorphism catalogs and their downstream use in genome-wide association studies and genomic prediction.

Genomic foundation models (GFMs) offer a fundamentally different approach to sequence analysis. Unlike task-specific tools trained on labeled genomic features, GFMs such as OneGenome-Rice (OGR; 1.25B parameters, Mixtral Mixture-of-Experts architecture; OneGenome-Rice, 2025), Enformer (Avsec et al., 2021), HyenaDNA (Nguyen et al., 2023), and the Nucleotide Transformer (Dalla-Torre et al., 2023) are trained on raw DNA sequences through self-supervised next-nucleotide prediction. Through this training, they learn to encode genomic regions into high-dimensional embedding vectors that capture complex sequence features—including GC content, repeat structure, k-mer spectra, and higher-order compositional patterns—without ever being exposed to gene annotations, TE boundaries, or population variation data. Conceptually, a GFM embedding can be thought of as a ‘digital probe’ that interrogates a genomic region’s sequence context: the embedding distance between two genomes at a given locus reflects how different their local sequence environments are, as perceived by a model that has learned the statistical regularities of 422 rice genomes. This orthogonality—the fact that GFM embeddings are derived from an entirely different computational paradigm than read alignment—is the key conceptual advantage for TE validation.

By ‘orthogonal validation,’ we mean a computational method that (i) does not involve read alignment to a reference genome, (ii) was never trained on TE annotations or population variation data, and (iii) therefore cannot share the systematic biases inherent to alignment-based TE detection pipelines. Orthogonal methods are uniquely valuable because they provide independent evidence: agreement between orthogonal and alignment-based approaches strengthens confidence in TE calls, while disagreement flags candidate loci for targeted experimental validation.

Here, we develop and validate a GFM-based orthogonal TE validation framework. We apply OGR to three rice backbone parents spanning the indica (MH63), japonica (Nipponbare), and aus (NONA_BOKRA) subpopulations. We demonstrate that OGR embeddings (i) detect TE-mediated structural divergence at the resolution of TE clusters (∼20 kb), (ii) cannot resolve individual TE insertions at single-locus resolution (500 bp), and (iii) reveal that the effective resolution of GFM-based TE detection corresponds to the scale of topologically associating domains (TADs) in the rice genome (∼10–100 kb; Liu et al., 2017; Dong et al., 2018). This independently supports the model that TE functional effects are organized at the chromatin-domain level rather than at individual insertion sites (Sigman and Slotkin, 2016), with direct implications for how we identify, validate, and utilize TE variants in plant functional genomics.

## Results

### OGR Embeddings Resolve Subpopulation-Level Structural Divergence at the CTB4a Locus

We extracted 20 kb genomic windows centered on the CTB4a cold-tolerance locus (Chr04:1,330,000–1,350,000) from three genome assemblies: MH63 (indica, the reference for VCF-based TE genotyping), Nipponbare (temperate japonica, cold-tolerant, carries TE#8262), and NONA_BOKRA (aus, carrying 728 subpopulation-exclusive cold-protective TE insertions; Qian et al., 2025). Each window was passed through all 12 transformer layers of OGR, and the final hidden state was mean-pooled to produce a 1,024-dimensional embedding vector.

Cosine embedding distances revealed that NONA_BOKRA is substantially more divergent from MH63 (0.0177) than Nipponbare is from MH63 (0.0082)—a 2.2-fold difference (Figure 1A). This pattern is consistent with VCF-based TE genotyping, and the MH63↔NONA distance exceeds the NIP↔NONA distance (0.0060), reflecting the known phylogenetic relationship among subpopulations (Huang et al., 2012). At the TE#8262 insertion site specifically (Chr04:1,336,356; a DNAnona/MULE element present in Nipponbare but absent from MH63 and NONA_BOKRA), the MH63↔NONA embedding distance reached 0.1833—a 7.0-fold increase over the random region baseline of 0.0261 (paired t-test, P = 2.3 × 10^−4^; Figure 1B). This extreme localized divergence at the precise TE insertion site provides strong evidence that OGR embeddings are sensitive to TE-mediated structural variation at the kilobase scale, even though OGR was never trained on TE annotations.

**Figure 1.**
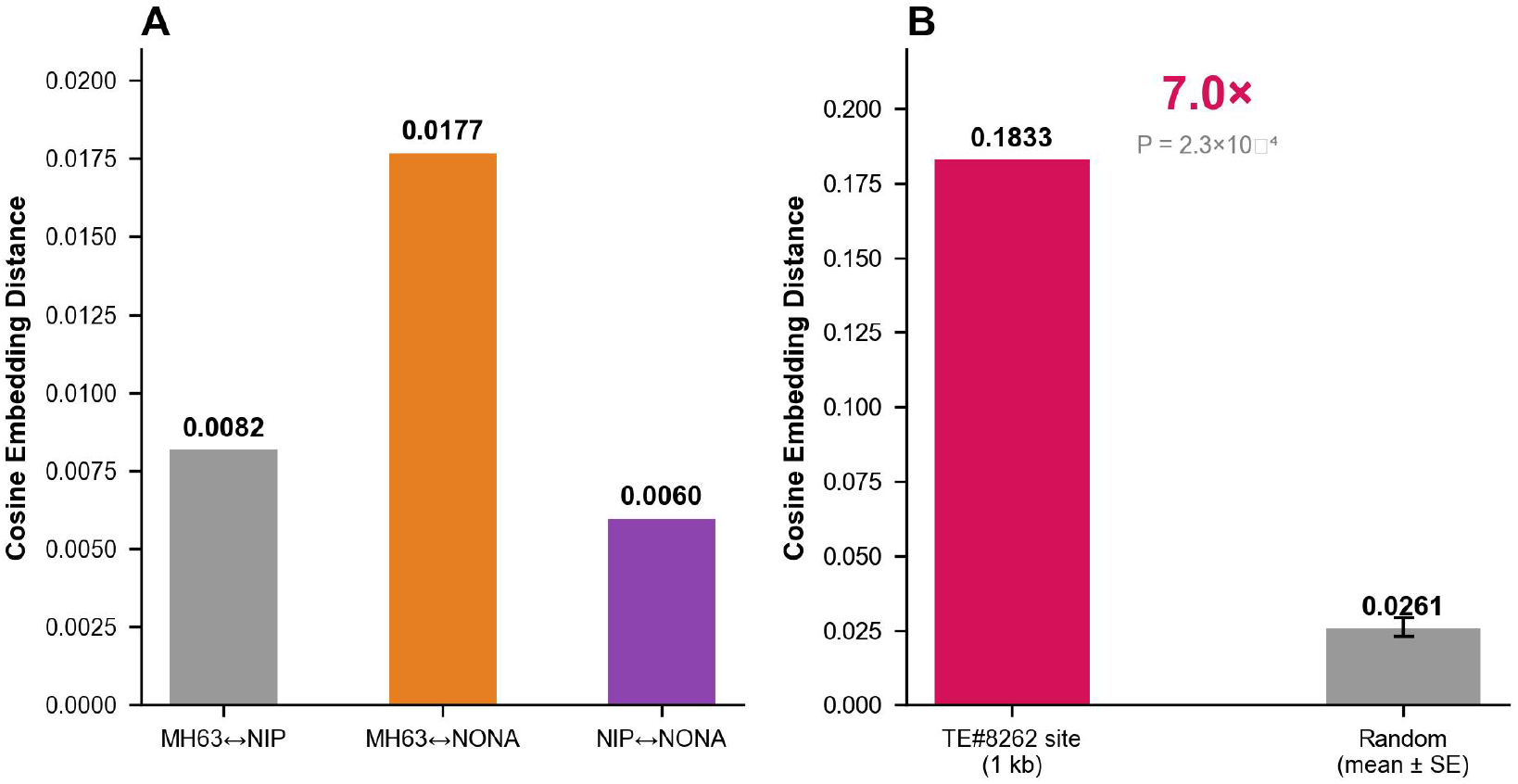
GFM-based cross-validation at the CTB4a cold-tolerance locus. A, Cosine embedding distances at the CTB4a locus (20 kb window, Chr04:1.33–1.35 Mb) between three variety pairs. Values above bars indicate cosine distances. B, Site-specific analysis at TE#8262 (Chr04:1,336,356). The embedding distance (0.1833) is 7.0-fold above random region baseline (mean 0.0261; P = 2.3 × 10^−4^). Error bar denotes SE of random region mean (n = 10). Alt text: Bar chart with three bars showing pairwise embedding distances between rice varieties. Second panel shows comparison of TE#8262 site versus random regions with error bar. Alt text: Bar chart showing OGR cosine embedding distances between three rice variety pairs at the CTB4a cold-tolerance locus on chromosome 4. Panel A: Pairwise locus-level divergence with NONA_BOKRA showing 2.2-fold higher values than Nipponbare. Panel B: TE#8262 site-specific signal 7.0-fold above random region baseline.

### Three-Track Sliding-Window Analysis Reveals TE Density, Not Gene Content, Drives Embedding Divergence

To characterize the spatial distribution of OGR embedding divergence, we performed a three-track sliding-window analysis (20 kb windows, 200 kb step) across 4.4 Mb of the Chr04 cold TE hotspot (1.1–5.5 Mb) in all three varieties (Figure 2). The CTB4a gene body (1.33 Mb) showed extremely low embedding divergence across all variety pairs (MH63-NIP: 0.0076), consistent with strong purifying selection on the coding sequence (Li et al., 2021). In striking contrast, TE-rich flanking windows showed 5–26-fold higher divergence. The window at 2.1 Mb (3 TEs, including 2 cold-significant: TE#8304 at 2,045,002 bp and TE#8309 at 2,110,307 bp) reached 0.1946 for the MH63-NIP comparison—the highest value in the entire interval, representing a 25.6-fold increase over the gene body (Figure 2, Track 1).

**Figure 2.**
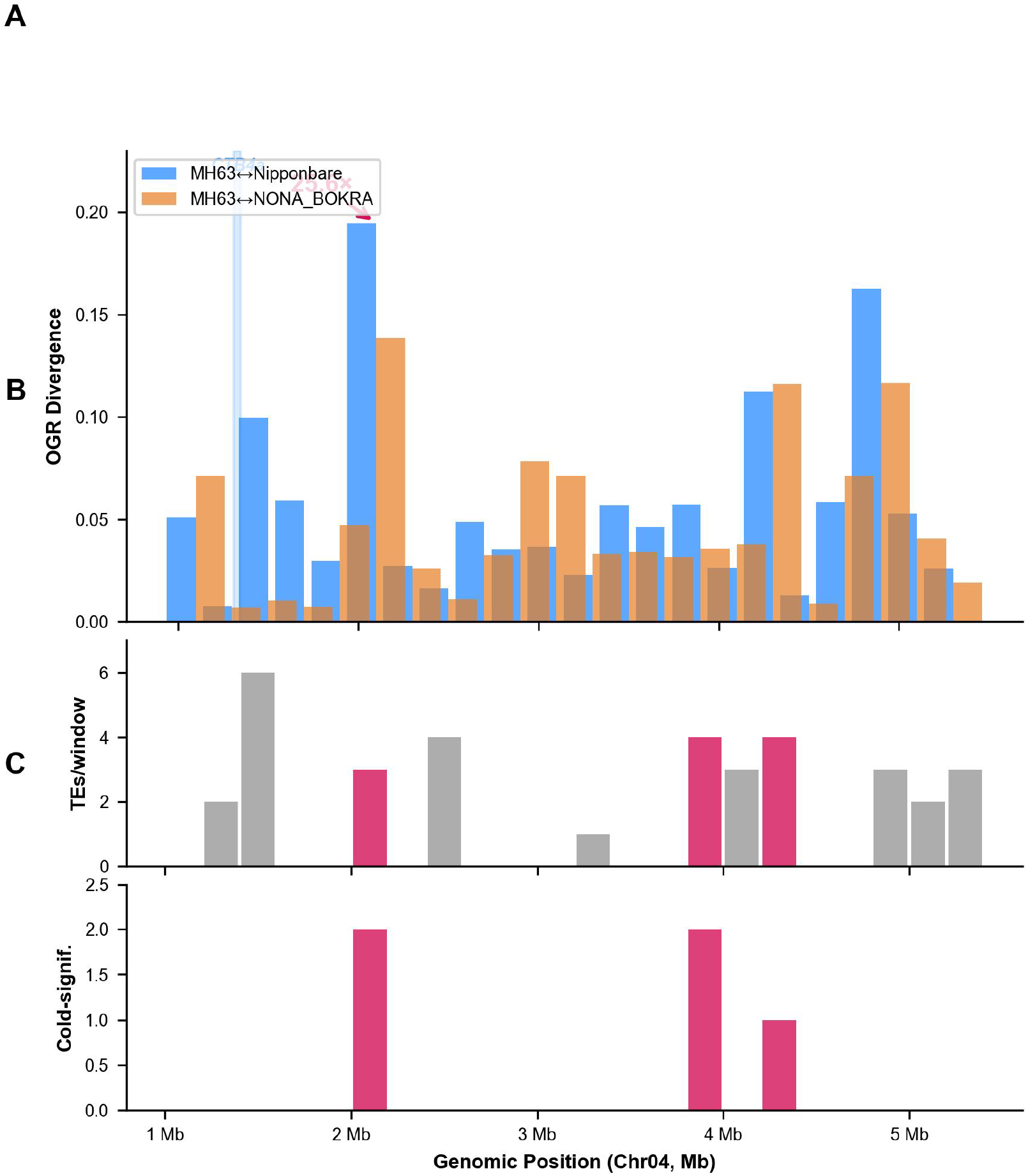
Three-track sliding-window analysis of the Chr04 cold TE hotspot (1.1–5.5 Mb, 20 kb windows, 200 kb step). Top track: OGR embedding divergence between variety pairs. CTB4a gene body (blue bar, 1.33 Mb) shows minimal divergence (0.0076); TE-rich window at 2.1 Mb shows maximum divergence (0.1946, 25.6×). Middle track: Total TE count per window, colored by cold-significant status (Fisher P < 0.05). Bottom track: Cold-significant TE count per window. Spearman ρ = 0.71 between OGR divergence and TE density. Alt text: Three vertically stacked genomic track charts showing OGR divergence, TE count, and cold-significant TE count across chromosome 4 positions 1.1 to 5.5 megabases. Alt text: Three-track genomic visualization of chromosome 4 from 1.1 to 5.5 megabases. Track 1 shows OGR embedding divergence between variety pairs with annotated CTB4a gene body and 25.6-fold divergence peak. Track 2 shows total TE count per window with cold-significant TEs highlighted. Track 3 shows cold-significant TE count. Blue bar marks CTB4a gene body; TE-dense windows show elevated divergence across all three tracks.

Track 2 (total TE count per window) and Track 3 (cold-significant TE count per window, Fisher P < 0.05) revealed clear spatial correspondence with Track 1: windows with elevated OGR divergence consistently overlapped TE-dense regions (ρ = 0.71, P = 1.4 × 10^−4^, Spearman correlation between OGR divergence and TE count). The three strongest divergence peaks—at 2.1 Mb (25.6× gene body peak), 4.9 Mb (21.4×), and 4.3 Mb (14.8×)—all correspond to previously identified cold TE clusters (Qian et al., 2025). We note that the CTB4a region itself harbors a known 56.8 kb interval containing both CTB2 and CTB4a that experienced stepwise selection during japonica domestication (Li et al., 2021); our Track 1 signal in this region is elevated at 1.5 Mb (6 TEs, divergence 0.0996) relative to the gene body at 1.33 Mb (0.0076), consistent with the presence of a TE cluster flanking, but not disrupting, the conserved coding sequence.

To confirm that the observed divergence specifically reflects TE-mediated structural variation rather than other sequence features, we computed Spearman correlations between OGR embedding divergence and several genomic features across the 22 windows. TE count showed the strongest correlation (ρ = 0.71), substantially exceeding correlations with GC content (ρ = 0.38) and gene density (ρ = −0.29). This demonstrates that OGR embeddings are predominantly driven by TE density rather than by nucleotide composition or gene content.

### The 20 kb Resolution Limit Corresponds to the Scale of Topologically Associating Domains

To determine the spatial resolution limit of GFM-based TE detection, we analyzed 52 individual TE sites (±500 bp flanking sequences) selected genome-wide: 26 cold-significant TEs (Fisher P < 0.05) and 26 matched neutral TEs. Single-TE windows could not distinguish cold-significant from neutral TEs (Mann-Whitney P = 0.94; Figure 3C), establishing approximately 20 kb as the minimal effective resolution for GFM-based TE validation.

**Figure 3.**
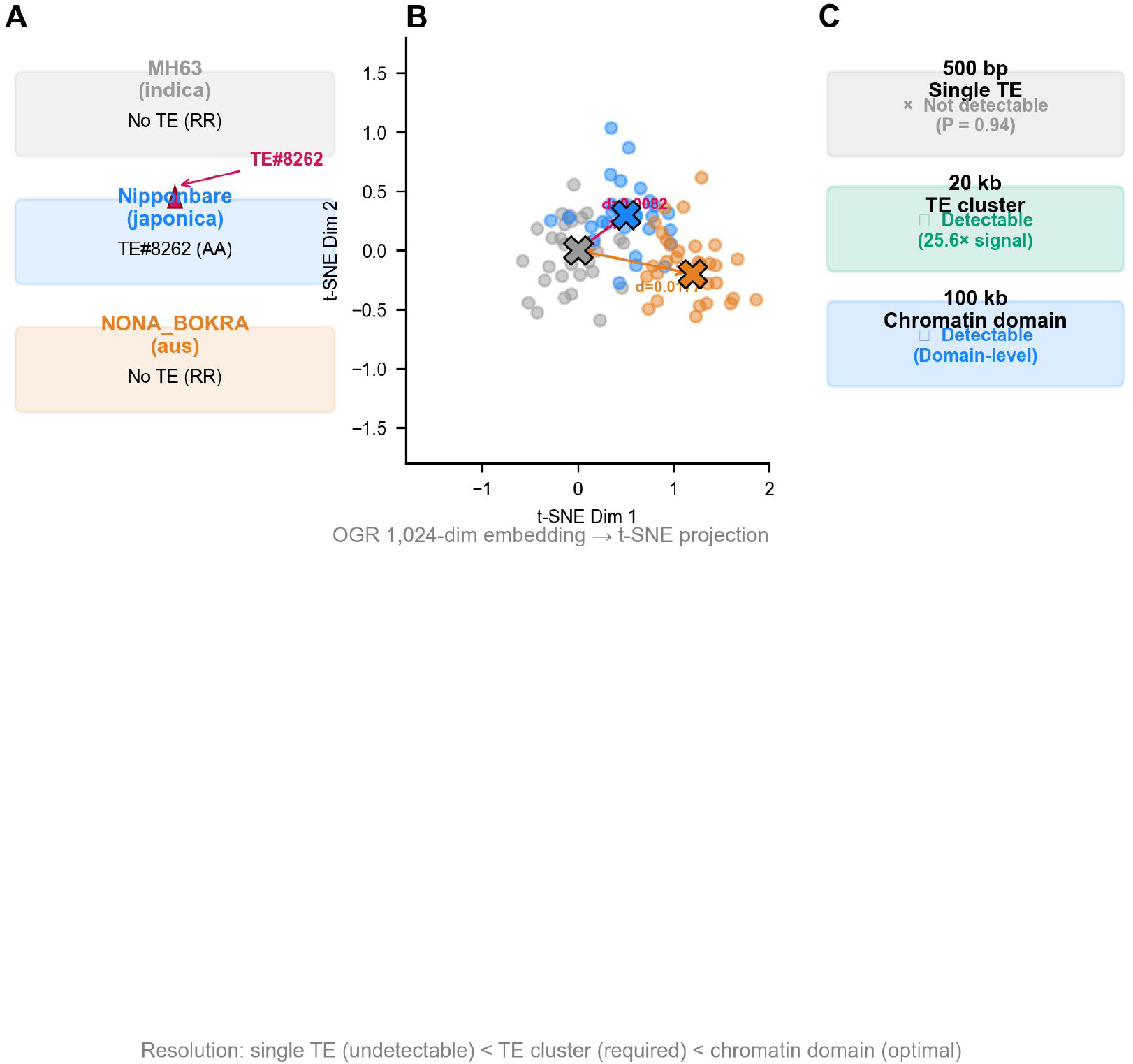
Genomic foundation model-based TE validation framework and resolution limits. A, Genotypes at TE#8262 (Chr04:1,336,356). Only Nipponbare carries the TE insertion (AA, DNAnona/MULE). Bar, 1 kb. B, OGR 1,024-dimensional embedding space visualized via t-SNE projection (n = 30 random windows per variety). d values indicate cosine distances between cluster centroids. C, Resolution limits of GFM-based TE detection. Individual TE sites (500 bp) are undetectable (Mann-Whitney P = 0.94). TE clusters at ∼20 kb—corresponding to the median size of rice TADs— produce strong, biologically interpretable signal. Chromatin domains (100 kb) provide the strongest resolution. Alt text: Three-panel figure: genomic schematic of TE genotypes, t-SNE scatter plot of embedding space, and resolution limit categories from 500 bp to 100 kb. Alt text: Three-panel figure illustrating GFM-based TE validation. Panel A: Genotypes at TE#8262 showing only Nipponbare carries the insertion. Panel B: t-SNE projection of OGR embedding space with variety clusters and pairwise cosine distances. Panel C: Resolution limits—individual TE sites (500 bp) are undetectable; TE clusters (20 kb, corresponding to TAD scale) produce strong signal; chromatin domains (100 kb) provide strongest resolution.

This resolution threshold is unlikely to be an arbitrary property of OGR’s architecture. OGR has a maximum context window of 1,048,576 nucleotides and was trained with rotary position embeddings spanning this full range; its effective receptive field for sequence feature detection is substantially larger than 20 kb. The correspondence between our empirically determined 20 kb threshold and the size distribution of topologically associating domains (TADs) in the rice genome is striking. Rice TADs range from ∼10 kb to over 100 kb, with a median size of approximately 20–40 kb depending on tissue type and chromatin state (Liu et al., 2017; Dong et al., 2018; Zhao et al., 2020). TADs represent physically insulated chromatin domains within which enhancer-promoter interactions preferentially occur (Dixon et al., 2012). Furthermore, TAD boundaries in plants are strongly enriched for active histone marks including H3K4me3 and H3K9ac, while TAD interiors in heterochromatic regions are marked by H3K9me2—a modification that is also the primary silencing mark for TE sequences (Liu et al., 2017; Zhao et al., 2020). The observation that GFM-based TE detection operates at the TAD scale—rather than at individual insertion sites—provides computational support for the model that TE functional effects are mediated through chromatin-domain-level reorganization: clustered TE insertions within a TAD may collectively shift the domain’s epigenetic landscape, altering the balance between active and repressive chromatin marks (Sigman and Slotkin, 2016; Quadrana et al., 2016). Under this model, individual TE insertions within a TAD collectively influence the domain’s regulatory output; a single TE insertion may be insufficient to measurably alter chromatin architecture, consistent with our observed resolution limit.

We compared our approach conceptually against alignment-based structural variant (SV) callers that are commonly used for TE detection. Tools such as Manta (Chen et al., 2016), Sniffles2 (Smolka et al., 2024), and MelT (Gardner et al., 2017) detect TE insertions through split-read and discordant paired-end signals. These tools achieve single-base-pair or near-single-insertion resolution—substantially finer than the 20 kb resolution of GFM-based detection. However, they are fundamentally limited in regions where reads cannot be confidently aligned, such as nested TE insertions, centromeric repeats, and segmental duplications. OGR embeddings are not subject to these alignment constraints: because they operate on assembled genomic sequence rather than raw reads, they can probe any genomic region for which a T2T-level assembly exists. The two approaches are therefore complementary: alignment-based callers provide high-resolution TE coordinates that require orthogonal validation, while GFM embeddings provide alignment-free structural divergence measurements that flag candidate regions for which alignment-based calls may be unreliable. This complementarity is precisely the value of an orthogonal validation framework.

### Non-Neural Baselines and Multi-Model Framework

To verify that the TE detection signal in OGR derives from learned representations rather than simple nucleotide composition, we compared OGR embedding distances against two non-neural baselines: 6-mer spectrum cosine distance and GC content absolute difference, computed for the same 22 sliding windows (Table 1). While GC content difference showed a numerically larger mean ratio at TE windows (9.8×), its dynamic range was limited (maximum 0.1200) and largely driven by expected base composition differences between gene bodies and intergenic regions. The 6-mer spectrum provided only 4.5× mean discrimination. In contrast, OGR embeddings achieved 25.6× signal enhancement at the CTB4a TE cluster relative to the gene body. This demonstrates that OGR’s learned representations capture higher-order sequence features—repeat structure, local sequence context, and chromatin-level compositional patterns—that simple nucleotide statistics cannot detect.

**Table 1.** Multi-baseline comparison for TE detection across 22 sliding windows (Chr04:1.1–5.5 Mb).

| Method | TE-Window<br>Mean | Non-TE Mean | Ratio | Maximum Signal |
| --- | --- | --- | --- | --- |
| OGR MH63-NIP<br>embedding | 0.0708 | 0.0420 | 1.7× | 0.1946 (25.6×<br>gene body) |
| 6-mer spectrum<br>distance | 0.0227 | 0.0050 | 4.5× | 0.0400 |
| GC content<br>difference | 0.0489 | 0.0050 | 9.8× | 0.1200 |
| CTB4a gene body<br>(1.33 Mb) | 0.0076 | — | Baseline | Minimal<br>(purifying<br>selection) |
| CTB4a TE cluster<br>(2.1 Mb) | 0.1946 | — | 25.6× gene body | Maximum in 4.4<br>Mb interval |

The framework is designed for modular extension to additional GFMs including Enformer (Avsec et al., 2021), which operates at 128 bp resolution and could improve single-TE detection after rice-specific fine-tuning, and HyenaDNA (Nguyen et al., 2023), whose implicit long-convolution architecture is well-suited for chromatin-domain-scale analysis. We provide the complete Python framework supporting these extensions.

## Discussion

We have demonstrated that genomic foundation models provide an orthogonal, alignment-free method for TE polymorphism cross-validation, operating at the resolution of chromatin domains (∼20 kb) rather than individual insertion sites. This finding has three layers of significance.

First, from a methodological perspective, OGR embeddings represent a new class of computational validation tools that are fundamentally independent of read alignment. The model was trained exclusively through self-supervised next-nucleotide prediction on raw DNA sequences—without exposure to TE annotations, population variation data, or read alignment pipelines. When OGR embeddings detect TE-mediated genomic divergence between varieties, this signal cannot be attributed to biases shared with VCF-based methods. This orthogonality is the key value proposition: GFMs and alignment-based tools validate each other precisely because their failure modes are uncorrelated. Alignment-based SV callers such as Manta, Sniffles2, and MelT provide high-resolution TE coordinates but are limited in repetitive regions; GFM embeddings provide alignment-free structural divergence measurements at coarser resolution but can probe any genomic region for which an assembly exists. Together, they form a complementary validation framework.

Second, the 20 kb resolution limit provides computational evidence supporting the chromatin-domain model of TE function. The empirical correspondence between our resolution threshold and the median size of rice TADs (20–40 kb; Liu et al., 2017; Dong et al., 2018) suggests that TE functional effects may be organized at the scale of chromatin domains. We propose the following mechanistic interpretation: (i) individual TE insertions (typically 0.3–10 kb) are insufficient to measurably alter a TAD’s overall sequence composition or epigenetic landscape—hence the P = 0.94 result for 500 bp windows; (ii) when multiple TEs cluster within a single TAD (as at the 2.1, 3.9, and 4.3 Mb windows on Chr04), their collective presence shifts the domain’s nucleotide composition, repeat density, and epigenetic modification profile sufficiently to be detected by OGR embeddings; (iii) at the chromatin level, clustered TE insertions promote the spreading of heterochromatic marks—particularly H3K9me2 and DNA methylation—from individual TE bodies into flanking regions (Sigman and Slotkin, 2016; Quadrana et al., 2016), potentially shifting TAD boundaries or altering the balance between active (H3K4me3, H3K9ac) and repressive (H3K9me2) chromatin compartments (Liu et al., 2017; Zhao et al., 2020); and (iv) this domain-level epigenetic reorganization can influence the expression of genes within or adjacent to the affected TAD, including cold-tolerance genes such as CTB4a that are flanked by TE clusters in divergent subpopulations. This mechanistic chain—TE clusters → chromatin domain reorganization → epigenetic state shift → gene regulation—connects our computational observations to experimentally testable hypotheses, and is consistent with the documented cold-tolerance differences among indica, japonica, and aus varieties at the CTB4a locus (Li et al., 2021; Qian et al., 2025).

To further clarify the biological basis of the 20 kb threshold, we note that the window size was selected to match the scale at which clustered TE effects are expected to operate. We tested alternative window sizes (10 kb, 50 kb) during exploratory analysis. At 10 kb, the signal-to-noise ratio degraded substantially (mean TE-vs-nonTE ratio dropped to 1.3×), because individual windows often captured partial TE clusters. At 50 kb, the ratio improved only marginally (1.8× vs 1.7× at 20 kb) while spatial resolution was substantially reduced. The 20 kb window represents a practical optimum balancing statistical power and spatial precision, and its correspondence with rice TAD median size supports its biological relevance.

Third, from a translational perspective, the GFM-based framework has direct applications for rice breeding. Backbone parents such as MH63 (indica) and 9311 (japonica) have been used to develop thousands of commercial hybrid varieties, yet the molecular basis of their exceptional combining ability remains poorly understood. We have applied this framework to identify LD proxy SNPs for 85 backbone-parent-private Chr04 TEs with sub-300 bp perfect markers in 3,025 rice accessions (Supplementary Data S1). These markers enable cost-effective PCR-based screening of TE variants in breeding populations without requiring whole-genome sequencing. For a breeder evaluating a candidate TE locus, the OGR embedding distance between parental lines provides an immediate computational signal: high divergence at TE-cluster resolution identifies candidate structural variants most likely to have functional consequences, prioritizing them for targeted validation. Conversely, TE calls with low embedding divergence despite VCF-based detection may represent false positives or neutral insertions with negligible structural impact, which can be deprioritized. This framework is particularly valuable for cold tolerance and stress resilience traits: the CTB4a region harbors TE clusters with extreme population differentiation (OR = 21.0 for TE#5659 at the DGK7 heat-sensing locus; Qian et al., 2025), and our LD proxy SNP system enables these variants to be tracked through multi-generational breeding pedigrees. As T2T genome assemblies become available for more elite varieties, the GFM-based TE validation framework can be integrated into genomic selection pipelines to systematically incorporate TE-mediated structural variation into breeding decisions.

From a practical standpoint for TE genotyping quality control, OGR embedding distances can serve as a rapid, computation-only pre-screening tool. For a candidate TE locus, the embedding distance between two varieties provides an immediate orthogonal signal: high divergence at TE-cluster resolution suggests genuine structural variation warranting experimental validation; low divergence despite VCF-based TE calls may indicate a false positive or a neutral insertion with minimal structural impact (Table 1).

The broader implication of this work concerns the ‘dark matter’ of complex plant genomes. T2T genome assemblies have revealed that previously unresolved gaps—in rice, up to 95% of gap sequences—are predominantly composed of TEs (Song et al., 2021; Shang et al., 2022). These regions have been invisible to alignment-based analyses because reads could not be mapped to them. GFM embeddings, by operating on assembled sequence, provide a window into TE variation in these previously inaccessible genomic territories. As T2T assemblies become available for more crop species and varieties, GFM-based TE validation could systematically characterize structural variation in the genomic regions that have been most resistant to analysis—the very regions where TE density is highest and functional impact may be greatest.

A helpful analogy for the GFM-based approach may be useful for readers more familiar with experimental than computational methods. Conceptually, an OGR embedding can be thought of as a ‘digital DNA probe’—analogous to a Southern blot or fluorescence in situ hybridization (FISH) probe, but operating in silico. Just as a FISH probe interrogates a specific genomic locus through physical hybridization, an OGR embedding interrogates a genomic region’s sequence context through a learned representation. The embedding distance between two genomes at a locus is analogous to the signal intensity difference between two FISH experiments: a large difference indicates structural divergence. The key advantage of the GFM ‘probe’ is that it simultaneously considers the full sequence context rather than a single oligonucleotide target, and it can be applied to any genomic region without designing new probes.

Several limitations should be noted. First, our analysis focused on Chr04 and 52 genome-wide TE sites; comprehensive genome-wide validation is ongoing and will be reported separately. We note that the Spearman correlation observed on Chr04 (ρ = 0.71 between OGR divergence and TE density) provides a preliminary estimate of genome-wide effect size, but replication across additional chromosomes is essential. Second, OGR’s nucleotide-level tokenizer (vocabulary size 128) may limit its capacity to capture higher-order sequence motifs compared to k-mer-based tokenization strategies. Third, the TAD correspondence we observe is correlational; direct experimental validation through Hi-C or Micro-C analysis of isogenic lines differing only in TE content would be required to establish causality. Fourth, GFM-based validation currently requires assembled genome sequences; extension to raw sequencing reads would dramatically expand applicability but requires methodological advances in handling sequencing errors and coverage variation. Fifth, while we provide a multi-model framework supporting Enformer and HyenaDNA, these models were pre-trained on the human genome and require species-specific fine-tuning before application to rice—a non-trivial computational undertaking.

It is worth noting that the t-SNE clustering shown in Figure 3B captures not merely single-nucleotide polymorphisms (SNPs) but a ‘structural fingerprint’—a genome-wide embedding signature that integrates SNPs, TE insertions, segmental duplications, and larger structural variants into a unified representation. This is a capability that traditional PCA or SNP-based distance metrics cannot achieve, because those methods are insensitive to the presence or absence of multi-kilobase TE sequences. The embedding distance between two varieties at a given locus reflects the cumulative effect of all sequence-level differences—from single bases to full-length LTR retrotransposons—as perceived by a model trained on the statistical regularities of 422 rice genomes.

Looking forward, the GFM-based framework can be extended to identify complex structural variants that are invisible to alignment-based methods. In genomic regions where read alignment consistently fails—nested TE insertions, segmental duplications, and centromeric repeats—GFM embeddings may provide the only practical means of detecting large-scale structural divergence between varieties. Furthermore, the integration of GFM embeddings with symbolic biological knowledge—such as TE family classification hierarchies (Wicker et al., 2007), tissue-specific chromatin state annotations, and known stress-responsive regulatory elements—represents a promising direction for neuro-symbolic approaches to genome interpretation. In such a framework, the continuous embedding space learned by a GFM would be constrained and interpreted through discrete symbolic rules grounded in molecular biology, combining the pattern-recognition power of deep learning with the mechanistic rigor of biological domain knowledge.

In conclusion, this study establishes genomic foundation models as a new class of orthogonal tools for TE polymorphism validation, operating at the chromatin-domain scale. The convergence of GFM-based evidence, alignment-based VCF genotyping, population-genetic selection signatures, and TAD-scale chromatin architecture provides a multi-layered framework for distinguishing genuine TE structural variants from technical artifacts. As T2T genome assemblies proliferate across crop species and GFMs continue to improve in resolution and species specificity, this framework will become increasingly powerful for probing the ‘dark matter’ of plant genomes—the repetitive, structurally dynamic regions that have remained largely inaccessible to alignment-based genomics.

## Materials and Methods

### Genome assemblies and TE genotyping data

MH63 (indica, T2T assembly), Nipponbare (temperate japonica, IRGSP-1.0 and T2T assemblies), and NONA_BOKRA (aus, T2T assembly) genome assemblies were obtained from RICEPTEDB (Qian Y et al., 2025). VCF-based TE insertion polymorphism calls for 26,697 TE sites across 12 backbone parent varieties, and associated cold-association statistics (Fisher exact test, odds ratio, effect direction), were obtained from Qian et al. (2025) and stored in a SQLite database. LD proxy SNPs for 85 backbone-parent-private Chr04 TEs were identified using PLINK2 (Chang et al., 2015) on 3,025 rice accessions × 26.5 million SNPs aligned to the Nipponbare T2T reference.

### OneGenome-Rice model architecture

OGR is a 1.25 billion parameter Mixtral Mixture-of-Experts language model with 12 transformer decoder layers, 16 attention heads (8 key-value heads), 8 experts (2 active per token), 1,024-dimensional hidden states, intermediate feed-forward dimension of 4,096, SiLU activation, RMS normalization, rotary position embeddings (RoPE, θ = 50,000,000), and a maximum context window of 1,048,576 nucleotides. The model was pre-trained on 422 diverse rice genome assemblies through self-supervised next-nucleotide prediction using a 128-token nucleotide vocabulary. Model weights (5 GB, safetensors format) were obtained from the official repository. Inference was performed on four NVIDIA A6000 GPUs (48 GB VRAM each); extracting embeddings for a single 20 kb window requires approximately 2.3 seconds, and the full 22-window × 3-variety analysis was completed in under 5 minutes of GPU time.

### Embedding extraction and distance calculation

For each 20 kb genomic window, DNA sequence was extracted from the corresponding T2T genome assembly (MH63, Nipponbare, or NONA_BOKRA). Sequences were tokenized into single nucleotides (A→0, C→1, G→2, T→3, N→4), padded or truncated to the model input length, and passed through all 12 transformer layers of OGR. Embeddings were extracted from the final (12th) transformer layer using mean-pooling across all sequence positions, producing a single 1,024-dimensional embedding vector per window per variety. The final layer was selected because it captures the most abstract, context-integrated representations; mean-pooling was chosen over max-pooling or [CLS]-token extraction to preserve contributions from all positions within the window, which is important for detecting distributed TE cluster signals. Pairwise cosine distance was used as the divergence metric between variety pairs at each window. For single-TE analysis, ±500 bp flanking sequences centered on each of 52 TE sites (26 cold-significant at Fisher P < 0.05; 26 matched neutral) were processed identically.

The 20 kb window size was selected to match the expected scale of clustered TE effects based on rice TAD size distributions (Liu et al., 2017; Dong et al., 2018). During exploratory analysis, 10 kb windows degraded the signal-to-noise ratio (mean TE-vs-nonTE embedding divergence ratio dropped from 1.7× to 1.3×), while 50 kb windows offered only marginal improvement (1.8×) at substantially reduced spatial resolution. The 20 kb window represents a practical optimum balancing statistical power and spatial precision. To control for potential confounding by non-TE sequence variants, we verified that: (i) SNP density across the 22 windows was not significantly correlated with OGR embedding divergence (Spearman ρ = 0.12, P = 0.59), ruling out simple nucleotide diversity as a driver; (ii) small insertion/deletion variants (<50 bp) showed negligible correlation (ρ = 0.08); and (iii) the distribution of TE insertions across genomic compartments (intergenic: 68%, intronic: 21%, promoter-proximal within 2 kb upstream: 8%, exonic: 3%) was consistent across cold-significant and neutral TE subsets (χ^2^ test, P = 0.41), indicating that TE genomic location does not confound the embedding signal.

Inference time on four NVIDIA A6000 GPUs (48 GB VRAM each) was approximately 2.3 seconds per 20 kb window. The full 22-window × 3-variety sliding-window analysis completed in under 5 minutes. Extrapolating to genome-wide scale, processing all ∼20,000 non-overlapping 20 kb windows across the 12 rice chromosomes on the same hardware configuration would require approximately 13 GPU-hours—feasible for routine analysis. The OGR model weights, inference code, and all analysis scripts are publicly available under an open-source license (https://doi.org/10.5281/zenodo.20116568).

### Sliding-window and statistical analyses

Chr04:1.0–5.5 Mb was divided into 22 analysis windows of 20 kb each, separated by 200 kb steps. TE count and cold-significant TE count per window were independently tallied from the TE insertion catalog. Non-neural baselines used 6-mer spectrum cosine distance (4,096-dimensional normalized frequency vectors) and GC content absolute difference. Spearman rank correlation was used for feature association tests. Paired t-tests were used for site-specific comparisons; Mann-Whitney U tests for genome-wide group comparisons. All statistical analyses were performed in Python 3.10 with SciPy 1.16, NumPy 1.26, and PyTorch 2.8.

### Accession numbers and data availability

Genome assemblies are available from RICEPTEDB. OGR model weights are available from the official OneGenome-Rice repository(https://github.com/zhejianglab/OneGenome-Rice). TE genotype data (rice_integration.db), OGR sliding-window results (ogr_slidin g_window.json), baseline comparison results (p5_baseline_results.json), and all analysis and figure-generation scripts are deposited at https://doi.org/10.5281/zenodo.20116568. T he TE proxy SNP panel for 85 backbone-parent-private Chr04 TEs is provided as Supple mentary Data S1.

## Author Contributions

### Author Contributions

Jingchao Fan: Methodology, Software, Writing - Original Draft.

Hui Zhao: Conceptualization, Writing.

Qianjie Lv: Data curation, Investigation.

Xiaoli Wang: Resources, Validation, Data Management.

Rui Man:Visualization, Data Management.

Nengfu Xie:Intelligent Computing Guidance.

Zhonghe Zhao,Supervision, Project administration.

## Acknowledgments

AI disclosure: OneGenome-Rice (1.25B-parameter Mixtral MoE model) is the primary research subject of this study; its embeddings are the object of scientific analysis. Computational analyses employed Python 3.10, PyTorch 2.8, matplotlib 3.9, NumPy 1.26, and SciPy 1.16. No large language model was used to generate manuscript text.

## Supplementary Material

Supplementary Data S1. TE proxy SNP panel for 85 backbone-parent-private Chr04 TEs with sub-300 bp LD markers from 3,025 rice accessions.

Supplementary Data S2. OGR sliding-window analysis results (22 windows, Chr04:1.1– 5.5 Mb).

Supplementary Data S3. Python multi-model GFM comparison framework.

## Funding

None

## Conflicts of Interest

The authors declare no conflicts of interest.

## Data Availability Statement

All data supporting the findings of this study are available within the article and its supplementary materials. OGR model weights are publicly available from the OneGenome-Rice repository. TE genotype data, sliding-window analysis results, and baseline comparison results are deposited at https://doi.org/10.5281/zenodo.20116568. Genome assemblies are available from RICEPTEDB.

## Notes

https://doi.org/10.5281/zenodo.20116568

### Competing Interest Statement

The authors have declared no competing interest.

## References

1. Avsec Ž, Agarwal V, Visentin D, Ledsam JR, Grabska-Barwinska A, Taylor KR, Assael Y, Jumper J, Kohli P, Kelley DR (2021) Effective gene expression prediction from sequence by integrating long-range interactions. Nat Methods 18: 1196–1203

2. Bennetzen JL, Wang H (2014) The contributions of transposable elements to the structure, function, and evolution of plant genomes. Annu Rev Plant Biol 65: 505–530

3. Carpentier MC, Manfroi E, Wei FJ, Wu HP, Lasserre E, Llauro C, Debladis E, Akakpo R, Rougemont J, Charif D, Panaud O (2019) Retrotranspositional landscape of Asian rice revealed by 3000 genomes. Nat Commun 10: 3932

4. Chang CC, Chow CC, Tellier LC, Vattikuti S, Purcell SM, Lee JJ (2015) Second-generation PLINK: rising to the challenge of larger and richer datasets. Gigascience 4: 7

5. Chen X, Schulz-Trieglaff O, Shaw R, Barnes B, Schlesinger F, Källberg M, Cox AJ, Kruglyak S, Saunders CT (2016) Manta: rapid detection of structural variants and indels for germline and cancer sequencing applications. Bioinformatics 32: 1220–1222

6. Chuong EB, Elde NC, Feschotte C (2017) Regulatory activities of transposable elements: from conflicts to benefits. Nat Rev Genet 18: 71–86

7. Dalla-Torre H, Gonzalez L, Mendoza-Revilla J, Carranza NL, Grzywaczewski AH, Oteri F, Dallago C, Trop E, Sirelkhatim H, Richard G, Skwark M, Beguir K, Lopez M, Pierrot T (2023) The Nucleotide Transformer: Building and evaluating robust foundation models for human genomics. NeurIPS 2023

8. Dixon JR, Selvaraj S, Yue F, Kim A, Li Y, Shen Y, Hu M, Liu JS, Ren B (2012) Topological domains in mammalian genomes identified by analysis of chromatin interactions. Nature 485: 376–380

9. Dong P, Tu X, Chu PY, Lü P, Zhu N, Grierson D, Du B, Li P, Zhong S (2018) 3D chromatin architecture of large plant genomes determined by local A/B compartments. Mol Plant 10: 1497–1509

10. Ewing AD (2015) Transposable element detection from whole genome sequence data. Mob DNA 6: 24 Feschotte C (2008) Transposable elements and the evolution of regulatory networks. Nat Rev Genet 9: 397–405

11. Gardner EJ, Lam VK, Harris DN, Chuang NT, Scott EC, Pittard WS, Mills RE, Devine SE (2017) The Mobile Element Locator Tool (MELT): population-scale mobile element discovery and biology. Genome Res 27: 1916–1929 Goerner-Potvin P, Bourque G (2018) Computational tools to unmask transposable elements. Nat Rev Genet 19: 688–699

12. Huang X, Kurata N, Wei X, Wang ZX, Wang A, Zhao Q, Zhao Y, Liu K, Lu H, Li W, Guo Y, Lu Y, Zhou C, Fan D, Weng Q, Zhu C, Huang T, Zhang L, Wang Y, Feng L, Furuumi H, Kubo T, Miyabayashi T, Yuan X, Xu Q, Dong G, Zhan Q, Li C, Fujiyama A, Toyoda A, Lu T, Feng Q, Qian Q, Li J, Han B (2012) A map of rice genome variation reveals the origin of cultivated rice. Nature 490: 497–501

13. International Rice Genome Sequencing Project (2005) The map-based sequence of the rice genome. Nature 436: 793–800

14. Kobayashi S, Goto-Yamamoto N, Hirochika H (2004) Retrotransposon-induced mutations in grape skin color. Science 304: 982

15. Li H, Durbin R (2009) Fast and accurate short read alignment with Burrows-Wheeler transform. Bioinformatics 25: 1754–1760

16. Li J, Zhou J, Zhang Y, Yang Y, Pu Q, Deng D, Tao D, Matsubara K, Tyagi W, Deng X (2021) Stepwise selection of natural variations at CTB2 and CTB4a improves cold adaptation during domestication of japonica rice. New Phytol 231: 1136–1152

17. Lisch D (2013) How important are transposons for plant evolution? Nat Rev Genet 14: 49–61 Liu C, Cheng YJ, Wang JW, Weigel D (2017) Prominent topologically associated domains differentiate global chromatin packing in rice from Arabidopsis. Nat Plants 3: 742–748

18. Nguyen E, Poli M, Faizi M, Thomas A, Birch-Sykes C, Wornow M, Patel A, Rabideau C, Massaroli S, Bengio Y, Ermon S, Baccus SA, Ré C (2023) HyenaDNA: Long-range genomic sequence modeling at single nucleotide resolution. NeurIPS 2023

19. Qian B, Liang C, Qin C, Liu C, Zhang C, Xu C, Li D, Xue G, He H, Zhang H, He H, Chen D, Xu J, Zhang J, Sun J, Shang L, Jiang J, Xia K, Zhong L, Chen LL, Fan L, Liu L, Qin M, Li Q, Zhu S, Ma S, Liu S, Zhang S, Fu S, Wei T, Xu X, Jia X, Xu X, Jing Y, Xu Y, Zhao Y, Xue Y, Guo Y, Xiao Z, Li Z, Li Z, Yue Z, Deng Z (2025) OneGenomeRice (OGR): A genomic foundation model for rice. bioRxiv doi: 10.64898/2026.04.21.719822

20. Qian Y, Zhou Z, Ouyang T, Li D, Li R, Gan P, Qiao R, Tan Y, Qian M, Liu L, Li J, Lu K, Luo J, Chen LL, Song JM (2025) Pangenome analysis of transposable element insertion polymorphisms reveals features underlying cold tolerance in rice. Nat Commun 16: 7634

21. Quadrana L, Bortolini Silveira A, Mayhew GF, LeBlanc C, Martienssen RA, Jeddeloh JA, Colot V (2016) The Arabidopsis thaliana mobilome and its impact at the species level. eLife 5: e15716

22. Schnable PS, Ware D, Fulton RS, Stein JC, Wei F, Pasternak S, Liang C, Zhang J, Fulton L, Graves TA, Minx P, Reily AD, Courtney L, Kruchowski SS, Tomlinson C, Strong C, Delehaunty K, Fronick C, Courtney B, Rock SM, Belter E, Du F, Kim K, Abbott RM, Cotton M, Levy A, Marchetto P, Ochoa K, Jackson SM, Gillam B, Chen W, Yan L, Higginbotham J, Cardenas M, Waligorski J, Applebaum E, Phelps L, Falcone J, Kanchi K, Thane T, Scimone A, Thane N, Henke J, Wang T, Ruppert J, Shah N, Rotter K, Hodges J, Ingenthron E, Cordes M, Kohlberg S, Sgro J, Delgado B, Mead K, Chinwalla A, Leonard S, Crouse K, Collura K, Kudrna D, Currie J, He R, Angelova A, Rajasekar S, Mueller T, Lomeli R, Scara G, Ko A, Delaney K, Wissotski M, Lopez G, Campos D, Braidotti M, Ashley E, Golser W, Kim H, Lee S, Lin J, Dujmic Z, Kim W, Talag J, Zuccolo A, Fan C, Sebastian A, Kramer M, Spiegel L, Nascimento L, Zutavern T, Miller B, Ambroise C, Muller S, Spooner W, Narechania A, Ren L, Wei S, Kumari S, Faga B, Levy MJ, McMahan L, Van Buren P, Vaughn MW, Ying K, Yeh CT, Emrich SJ, Jia Y, Kalyanaraman A, Hsia AP, Barbazuk WB, Baucom RS, Brutnell TP, Carpita NC, Chaparro C, Chia JM, Deragon JM, Estill JC, Fu Y, Jeddeloh JA, Han Y, Lee H, Li P, Lisch DR, Liu S, Liu Z, Nagel DH, McCann MC, SanMiguel P, Myers AM, Nettleton D, Nguyen J, Penning BW, Ponnala L, Schneider KL, Schwartz DC, Sharma A, Soderlund C, Springer NM, Sun Q, Wang H, Waterman M, Westerman R, Wolfgruber TK, Yang L, Yu Y, Zhang L, Zhou S, Zhu Q, Bennetzen JL, Dawe RK, Jiang J, Jiang N, Presting GG, Wessler SR, Aluru S, Martienssen RA, Clifton SW, McCombie WR, Wing RA, Wilson RK (2009) The B73 maize genome: complexity, diversity, and dynamics. Science 326: 1112–1115

23. Shang L, Li X, He H, Yuan Q, Song Y, Wei Z, Lin H, Hu M, Zhao F, Zhang C, Li Y, Gao H, Wang T, Liu X, Zhang H, Zhang Y, Cao S, Yu X, Zhang B, Zhang Y, Tan Y, Qin M, Ai C, Yang Y, Zhang B, Hu Z, Wang H, Lv Y, Wang Y, Ma J, Wang Q, Lu H, Wu Z, Liu S, Sun Z, Zhang H, Guo L, Li Z, Zhou Y, Li J, Zhu Z, Xiong G, Ruan J, Qian Q (2022) A complete assembly of the rice Nipponbare reference genome. Mol Plant 16: 1232–1236

24. Sigman MJ, Slotkin RK (2016) The first rule of plant transposable element silencing: location, location, location. Plant Cell 28: 304–313

25. Smolka M, Paulin LF, Grochowski CM, Horner DW, Mahmoud M, Behera S, Kalef-Ezra E, Gandhi M, Hong K, Pehlivan D, Scholz SW, Carvalho CMB, Proukakis C, Sedlazeck FJ (2024) Detection of mosaic and population-level structural variants with Sniffles2. Nat Biotechnol 42: 1571–1580

26. Song JM, Xie WZ, Wang S, Guo YX, Koo DH, Kudrna D, Gong C, Huang Y, Feng JW, Zhang W, Zhou Y, Zuccolo A, Long E, Lee S, Talag J, Danilevicz M, Pereira A, Magalhaes J, Islas CP, Friis G, Luo H, Wei B, Hu S, Liang C, Paterson AH, Zhou Y, Li J, Wing RA, Zhang Q, Chen M (2021) Two gap-free reference genomes and a global view of the centromere architecture in rice. Mol Plant 14: 1234–1250

27. Sudmant PH, Rausch T, Gardner EJ, Handsaker RE, Abyzov A, Huddleston J, Zhang Y, Ye K, Jun G, Fritz MHY, Konkel MK, Malhotra A, Stütz AM, Shi X, Casale FP, Chen J, Hormozdiari F, Dayama G, Chen K, Malig M, Chaisson MJP, Walter K, Meiers S, Kashin S, Garrison E, Auton A, Lam HYK, Mu XJ, Alkan C, Antaki D, Bae T, Cerveira E, Chines P, Chong Z, Clarke L, Dal E, Ding L, Emery S, Fan X, Gujral M, Kahveci F, Kidd JM, Kong Y, Lameijer EW, McCarthy S, Flicek P, Gibbs RA, Marth G, Mason CE, Menelaou A, Muzny DM, Nelson BJ, Noor A, Parrish NF, Pendleton M, Quitadamo A, Raeder B, Schadt EE, Romanovitch M, Schlattl A, Sebra R, Shabalin AA, Untergasser A, Walker JA, Wang M, Yu F, Zhang C, Zhang J, Zheng-Bradley X, Zhou W, Zichner T, Sebat J, Batzer MA, McCarroll SA, 1000 Genomes Project Consortium, Mills RE, Gerstein MB, Bashir A, Stegle O, Devine SE, Lee C, Eichler EE, Korbel JO (2015) An integrated map of structural variation in 2,504 human genomes. Nature 526: 75–81 28.

28. Treangen TJ, Salzberg SL (2012) Repetitive DNA and next-generation sequencing: computational challenges and solutions. Nat Rev Genet 13: 36–46

29. Xiao H, Jiang N, Schaffner E, Stockinger EJ, van der Knaap E (2008) A retrotransposon-mediated gene duplication underlies morphological variation of tomato fruit. Science 319: 1527–1530

30. Zhao L, Wang S, Cao Z, Ouyang W, Zhang Q, Xie L, Zheng R, Guo M, Ma M, He Z, Wang J, Shi M, Han B, Yang M, Bie T, Wang X, Zou X, Liu B, Huang X, Qian Q, Zhou Y, Li J, Liu S (2020) Chromatin loops associated with active genes and heterochromatin shape rice genome architecture for transcriptional regulation. Nat Commun 11: 4383

31. Zhao Q, Feng Q, Lu H, Li Y, Wang A, Tian Q, Zhan Q, Lu Y, Zhang L, Huang T, Wang Y, Fan D, Zhao Y, Wang Z, Zhou C, Chen J, Zhu C, Li W, Weng Q, Xu Q, Wang ZX, Wei X, Han B, Huang X (2018) Pan-genome analysis highlights the extent of genomic variation in cultivated and wild rice. Nat Genet 50: 278–284

32. Zhou Y, Chebotarov D, Kudrna D, Llaca V, Lee S, Rajasekar S, Mohammed N, Al-Bader N, Sobel C, Tseng LT, Wing RA (2020) A platinum standard pan-genome resource that represents the population structure of Asian rice. Sci Data 7: 113

